# Intergenerational Effects of Early Life Starvation on Life-History, Consumption, and Transcriptome of a Holometabolous Insect

**DOI:** 10.1101/2021.03.11.434967

**Authors:** Sarah Catherine Paul, Pragya Singh, Alice B. Dennis, Caroline Müller

## Abstract

Intergenerational effects, also known as parental effects in which the offspring phenotype is influenced by the parental phenotype, can occur in response to factors that occur not only in early but also in late parental life. However, little is known about how these parental life stage-specific environments interact with each other and with the offspring environment to influence offspring phenotype, particularly in organisms that realize distinct niches across ontogeny. We examined the effects of parental larval starvation and adult reproductive environment on offspring traits under matching or mismatching offspring larval starvation conditions using the holometabolous, haplo-diploid insect *Athalia rosae* (turnip sawfly). We show that the parental larval starvation treatment had trait-dependent intergenerational effects on both life-history and consumption traits of offspring larvae, partly in interaction with offspring conditions and sex, while there was no significant effect of parental adult reproductive environment. In addition, while offspring larval starvation led to numerous gene- and pathway-level expression differences, parental larval starvation impacted fewer genes and only the ribosomal pathway. Our findings reveal that parental starvation evokes complex intergenerational effects on offspring life-history traits, consumption patterns as well as gene expression, although the effects are less pronounced than those of offspring starvation.

## Introduction

Intergenerational effects, more commonly known as parental effects, are defined as causal influences of parental phenotype on offspring phenotype, likely mediated by non-DNA sequence-based inheritance (Wolf and Wade 2009; Perez and Lehner 2019). Together with transgenerational effects, which occur across several generations, they play a key role in the ecology and evolution of organisms (Badyaev and Uller 2009) and impact the responses of individuals to changing environmental conditions (Sánchez-Tójar et al. 2020). One important environmental factor that can rapidly change during an organism’s lifetime and across generations is food availability. Periods of starvation are commonly encountered by insects in the wild (Jiang et al. 2019) and, when experienced early in life, often have long-lasting consequences on an individual’s phenotype (Miyatake 2001; Wang et al. 2016; McCue et al. 2017) and the phenotype of the offspring (Saastamoinen et al. 2013; McCue et al. 2017; Paul et al. 2019). However, it is also becoming increasingly apparent that to fully elucidate the adaptive nature of intergenerational effects, studies should encompass different phases in development of the parental life cycle and how such life stage-specific experience affects offspring phenotype (English and Barreaux 2020).

The conditions experienced particularly during two phases of parental life, early life and later life during mating and reproduction, can have lasting, sometimes irreversible impacts on development trajectories of offspring (Monaghan 2008; Burton and Metcalfe 2014; Taborsky 2017). A favorable environment experienced in early life may result in better conditioned offspring (van de Pol et al. 2006) that can better cope with stress (Franzke and Reinhold 2013). This is often called silver spoon or carry-over effect (Monaghan 2008). Negative effects of a poor start in life might also be passed on, resulting in offspring less able to cope with stressful conditions (Naguib et al. 2006). Alternatively, such stressed individuals may produce offspring that are buffered against stressors, for example, through parental provisioning (Valtonen et al. 2012; Pilakouta et al. 2015; Hibshman et al. 2016). Finally, regardless of how favorable conditions are, it may be most important that they match between parents and offspring. Evidence for such predictive adaptive effects has been found in several species (Raveh et al. 2016; Le Roy et al. 2017), but is weak in others (Uller et al. 2013). Each of these trajectories of intergenerational effects are not mutually exclusive. For example, offspring in mismatched environments may do relatively better if their parents were of high quality, due to silver spoon effects (Engqvist and Reinhold 2016). Furthermore changes in parental investment based on mate cues (Cunningham and Russell 2000; Cornwallis and O’Connor 2009) may potentially counteract or augment the effects of the parental early life environment on offspring phenotype. As niches often shift across an individual’s lifetime, both parental early and later life environments and their interaction must be considered as factors that can influence offspring phenotypes.

With discrete phases during development, insects, particularly holometabolous insects, present ideal organisms in which to investigate how environmental cues experienced during early life but also during adulthood (e.g. mating) may interact with offspring environment, to influence individual offspring phenotypes (English and Barreaux 2020). Periods of larval starvation are known to influence developmental trajectories with knock-on effects on adult size, reproductive success and adult starvation resistance (Boggs and Niitepõld 2016; Wang et al. 2016). Individuals may respond to periods of starvation with increased food uptake (Regalado et al. 2017) as well as compensatory growth (i.e. a faster growth rate) or catch-up growth (i.e. attaining a minimum size by prolonging the larval development) (Hector and Nakagawa 2012). Moreover, starvation leads to substantial shifts in gene expression (Moskalev et al. 2015; Jiang et al. 2019; Etebari et al. 2020; Farahani et al. 2020). However, little is known about how long these effects may persist (McCue et al. 2017) and to what extent parental starvation may affect gene expression of offspring experiencing matching or mismatching conditions.

In the present study, we investigated the effects of parental larval starvation and parental reproductive environment on offspring (i.e. intergenerational effects), under matching or mismatching larval starvation conditions, using the turnip sawfly, *Athalia rosae* (Hymenoptera: Tenthredinidae). The larvae feed on leaves of various species of Brassicaceae, including crops, and can readily experience periods of starvation when their hosts are overexploited (Riggert 1939). The adults are nectar-feeding and in addition collect neo-clerodanoid-like compounds (hereafter called ‘clerodanoids’) from non-Brassicaceae plants. Clerodanoid-uptake improves their mating probability (Amano et al. 1999) and affects interactions between and within sexes (preprint Paul et al. 2021; preprint Paul and Müller 2021). This behavior is thus an important aspect of the adult life-history, but potential influences on offspring life-history traits have, to our knowledge, not yet been studied. We measured key life-history traits (i.e., developmental time and adult body mass) to assess the combined and interactive effects of parental and offspring larval resource availability and parental clerodanoid exposure on the offspring. Moreover, we measured effects of parental and offspring larval starvation on consumption and gene expression of offspring. We predicted that 1) there are stronger intragenerational than intergenerational effects of larval starvation treatment on all parameters measured; 2) matching conditions between parental and offspring starvation treatment are beneficial, while a mismatch leads to a reduced performance, with silver spoon or parental buffering effects potentially augmenting or dampening any mismatch effects; 3) clerodanoid exposure increases parental investment in offspring due to an enhanced mate attractiveness, leading to positive effects for the offspring (in terms of a faster development and higher body mass).

## Materials and Methods

### Set-Up of Insect Rearing and Plant Cultivation

Adults of *A. rosae* (F0) were collected in May 2019 at two locations (population A: 52°02’48.0”N 8°29’17.7”E, population B: 52°03’54.9”N 8°32’22.2”E). These individuals were reared for further two generations to reduce the impact of parental and grand-parental effects using a desing that minimized inbreeding (for details of breeding design see S1) White mustard (*Sinapis alba*) was provided for oviposition and Chinese cabbage (*Brassica rapa* var. *pekinensis*) as food plant. Adults of the F2 generation were kept individually in Petri dishes and provided with a honey:water mixture (1:50). *Athalia rosae* is haplodiploid, i.e. virgin females produce male offspring (Naito and Suzuki 1991). To increase the likelihood of gaining similar numbers of females and males, mated as well as virgin females were placed individually into boxes (25 x 15 x 10 cm). They were supplied with a middle-aged leaf of non-flowering cabbage plants for oviposition and a honey:water mixture, which was replenished daily. Females were removed from the boxes after one week and their offspring used to set up the experimental generations. Experimental rearing and consumption assays were carried out in a climate chamber (20 °C:16 °C, 16 h: 8 h light:dark, 70% r.h.).

Plants of *S. alba* and *B. rapa* were grown from seeds (Kiepenkerl, Bruno Nebelung GmbH, Konken, Germany) in a greenhouse (20 °C, 16 h: 8 h light:dark, 70% r.h.) and a climate chamber (20 °C, 16 h: 8 h light:dark, 70% r.h.). Plants of *Ajuga reptans* used for clerodanoid supply were grown from seeds (RHS Enterprise Ltd, London, UK) in the greenhouse and transferred outside in late spring once about 2 months old. Middle-aged leaves of plants that were about 8 month old were offered to adults.

### Experimental Overview and Measurements of Life-History Data

We conducted a fully factorial experiment where we manipulated the parental larval environment (parental larval starvation), the parental reproductive environment (exposure to clerodanoids) and the offspring environment (offspring larval starvation) (Fig. 1). The experimental generations were reared to test the effects of parental and offspring larval starvation, and their interaction with the adult reproductive environment on the offspring phenotype. On the day of hatching, larvae of the parental generation were individually placed into Petri dishes (5.5 cm diameter) with moistened filter paper and *B. rapa* leaf discs cut from 7-10 week old plants, which were replaced daily. Per maternal line, larvae were split equally between one of two larval starvation treatments, no starvation (N) or starvation (S). For starvation, individuals were starved twice for 24 h, first the day after moulting into 2^nd^ instar and second on the day of moulting into 4^th^ instar to minimize early mortality whilst mimicking the food deprivation larvae may experience when occurring in high densities (Riggert 1939). Larval instars were tracked by checking daily for the presence of exuviae.

**Figure 1:**
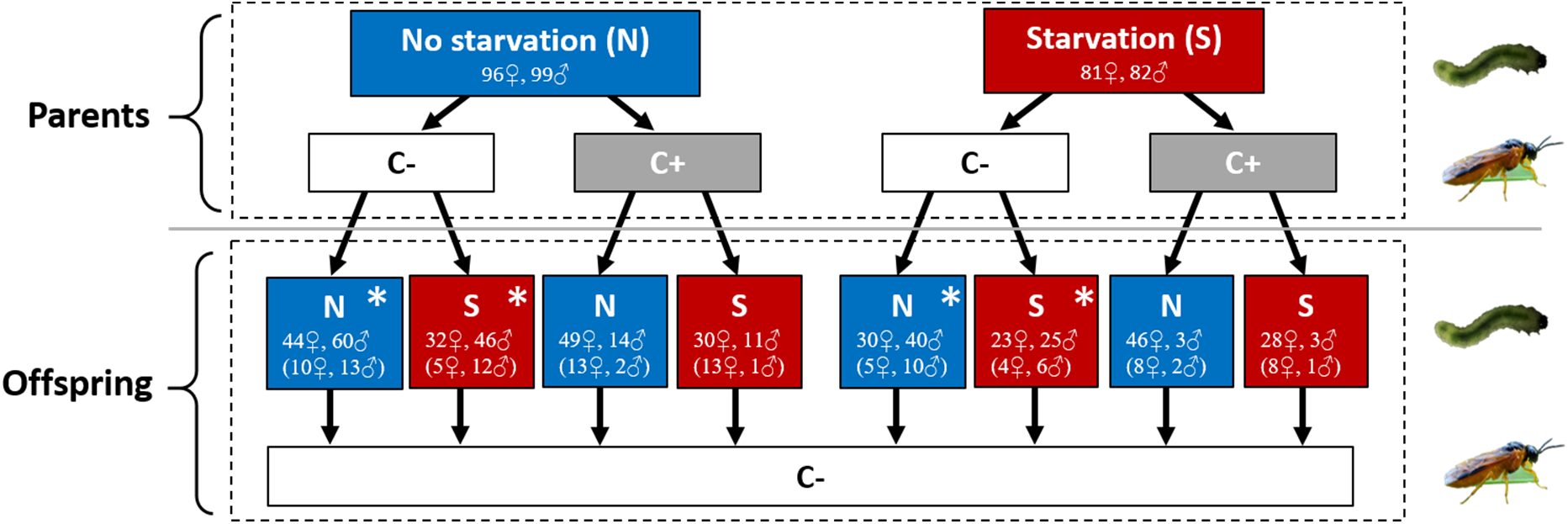
Design of experimental treatments with *Athalia rosae;* * indicates individuals taken for RNASeq analysis. Sample sizes split by sex are given in boxes. Numbers in brackets refer to sample sizes for consumption assay (please note that individuals were pooled across C+ and C-treatments). (For the full breeding design see S1.)

Eonymphs were placed in soil for pupation. Adults were kept individually in Petri dishes and provided with honey:water mixture. From the parental generation, pairs of non-sib females and males reared under the same larval starvation treatments were assigned to one of two reproductive environment treatments, where both parents either had clerodanoids (C+) or did not (C-). C+ individuals were exposed to a leaf section (1 cm^2^) of *A. reptans* for 48 h prior to mating, giving individuals time to take up clerodanoids (preprint Paul et al. 2021). Mated females (2-9 days old) from each of these four treatments (NC-, NC+, SC-, SC+) and C-virgin females (NC-, SC-) were then placed in individual breeding boxes. Their offspring were distributed to offspring starvation treatments (N or S) that matched the parental starvation treatment or differed from it (mismatch) (Fig. 1). In both generations, larval, pupal, and total development time (from larva to adult) as well as the adult body mass at the day of emergence (Sartorius AZ64, M-POWER Series Analytical Balance, Göttingen, Germany) were recorded for each individual. In total 358 larvae of the parental generation (177 females, 181 males) and 484 larvae of the offspring generation (282 females, 202 males) reached adulthood, out of the 607 and 688 larvae, respectively, that were reared in each generation.

### Consumption Assays

To test effects of parental and offspring larval starvation experience on offspring consumption, assays were performed with larvae of the offspring generation at the start of the 3^rd^ instar (directly after the first starvation event), measuring the relative growth rate, relative consumption rate, and efficiency of conversion of ingested food. The 3^rd^ instar was chosen to test the consumption not directly after a starvation event to avoid any potential interference with physiological changes directly induced by the starvation. Each larva was weighed at the beginning of the consumption assays (= initial body mass) (ME36S, accuracy 0.001 mg; Sartorius, Göttingen, Germany) and provided with four fresh discs cut from middle-aged leaves (surface area of 230.87 mm^2^ per disc) on moistened filter paper. After 24 hours, larvae were weighed again (= final body mass) and the leaf disc remains scanned (Samsung SAMS M3375FD, resolution 640 x 480). In that period, none of the leaf discs showed signs of wilting. The total area of leaf consumed (mm^2^) was then calculated as the difference between the average leaf area and the remaining leaf area. Leaf discs somewhat differ in mass but mass is likely more affected by different water contents than surface area.

### Statistical Analyses

All data were analyzed using R 4.0.2 (2020-06-22). We set *α* = 0.05 for all tests and checked model residuals for normality and variance homogeneity. All linear mixed effects models were run in lme4 using maximum likelihood. Stepwise backwards deletion using Chi^2^ ratio tests (package:MASS; version 7.3-53.1) for the life history traits and, due to the much smaller sample sizes (Luke 2017), conditional *F*-tests with *df* correction using Satterthwaite method (package lmerTest; version 3.1-3; Kuznetsova et al. 2017) for the consumption analyses were employed to reach the minimum adequate model (Crawley 2012). Posthoc analyses were carried out using the package ‘multcomp’ (version 1.4-13; Hothorn et al. 2008). Post data entry, raw data were visually inspected thrice, all variables plotted and outliers and possible anomalies in the data (e.g. strings of similar values) interrogated (package:pointblank; version 0.6.0). Intragenerational effects of starvation on life-history traits of individuals of the parental generation were tested as described in S1.

In *A. rosae*, usually 6 instars for female and 5 instars for males are found (Sawa et al. 1989). Due to observations made during the experiment (no *a priori* hypothesis), we tested in the offspring generation whether the likelihood of an additional larval instar (7 for females and 6 for males) differed based on offspring larval starvation treatment (independent of parental treatments) using a binomial generalized linear mixed model (package: lme4), where the predictor was offspring starvation treatment and parental pair was included as a random effect. The effects of the parental larval starvation treatment, parental clerodanoid exposure, offspring starvation treatment and their interaction on larval, pupal and total developmental time as well as adult mass of offspring individuals were assessed in separate linear mixed effects models (lmm), with parental pair included as a random effect (controlling for non-independence of sibling larvae). Female and male data were analyzed separately to enable model convergence.

Relative growth rate, relative consumption rate, and food conversion efficiency of larvae of the offspring generation were analyzed using lmms. We excluded the parental clerodanoid exposure as a predictor variable from the consumption assay analysis due to the low number of individuals in certain treatments (Fig. 1) and analyzed male and female data separately as above. To assess variation in relative growth rate, the change in larval body mass [final mass — initial mass] was used as the response variable and initial larval body mass, parental starvation treatment, offspring starvation treatment, and the interactions between all the predictors. To assess relative consumption rate, we used the total area of consumed leaf material as the response variable and initial larval body mass, parental starvation treatment, offspring starvation treatment, and both their three-way and two-way interactions as the predictors. Finally, for food conversion efficiency the change in larval body mass was taken as the response variable and the total area of consumed leaf material, parental starvation treatment, offspring starvation treatment, and the interactions between all the predictors. Parental pair was included in all three models as a random effect.

### Sample Collection and Sequencing

A total of 24 male larvae (4^th^ instar, 9 d old), comprising six biological replicates per treatment level (parent/offspring treatment: N/N, N/S, S/N, S/S, all C-, Fig. 2), were collected to investigate the effects of parental and offspring starvation treatment on gene expression in larvae of the offspring generation. Individuals were chosen in a way that maximized the equal spread of siblings across treatments and frozen at −80 °C prior to extraction. RNA was extracted with an Invitrogen PureLink™ RNA Mini Kit (ThermoFisher Scientific, Germany), including a DNase step (innuPREP DNase I Kit, analyticJena, Jena, Germany). RNA quality was assessed on a bioanalyzer 2100 (Agilent, CA, United States) and Xpose (PLT SCIENTIFIC SDN. BHD, Malaysia). Library preparation (Ribo-Zero for rRNA removal) and sequencing (NovaSeq6000 and S4 Flowcells, Illumina, CA, United States) were provided by Novogene (Cambridge, UK). Sequence quality before and after trimming was assessed using FastQC (v. 0.11.9; Andrews 2010). After short (< 75 bp), low quality (Q < 4, 25 bp sliding window), and adapter sequences were removed using Trimmomatic (Bolger et al. 2014), more than 98% of reads remained.

**Figure 2:**
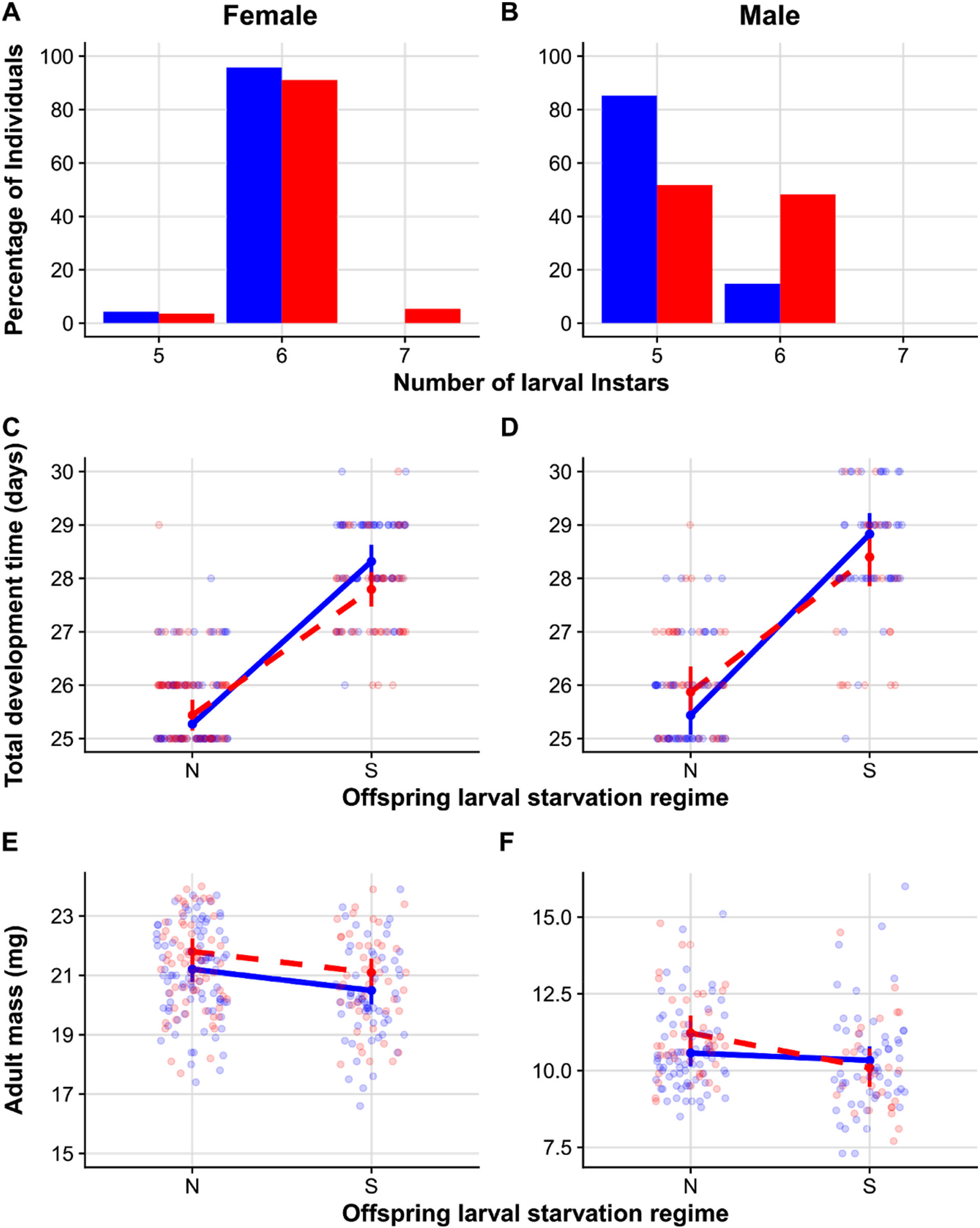
Number of larval instars in in dependence of offspring starvation treatment (blue: non starved, red: starved) in percentage (A, B) and influence of offspring larval starvation treatments (N = no starvation, S = starvation) and parental larval starvation treatments (blue solid line = no starvation, red dashed line = starvation) on offspring total development time (C, D) and adult body mass (E, F) of *Athalia rosae*. Data are plotted separately for females (A, C, E) and males (B, D, F); individuals were sexed on emergence. Points (C-F) are model predictions with associated confidence intervals and colors of points and lines correspond to parental starvation treatment. Raw data (C-F) are plotted in transparent colors in the background (blue circles - no parental starvation, red circles - parental starvation).

### Differential Expression Analysis

Cleaned reads were mapped to the annotated genome of *A. rosae*, version AROS v.2.0 (GCA_000344095.2) with RSEM v1.3.1 (Li and Dewey 2011), which implemented mapping with STAR v2.7.1a (Dobin et al. 2013). Analysis of differential gene expression was conducted with DESeq2 (version 1.28.1; Love et al. 2014). The results of mapping with RSEM were passed to DESeq2 for gene-level analyses using Tximport (version 1.16.1; Soneson et al. 2015). Prior to analysis, genes with zero counts in all samples and those with low counts (< 10) in less than a quarter of samples (6) were excluded. Model fitting was assessed by plotting dispersion estimates of individual gene models and outlier samples were inspected using principle component analysis of all expressed genes and pairwise-distance matrices between samples. Expression was modelled based on the entire dataset, with the four levels representing the combination of parental.offspring starvation treatments: N.N, N.S, S.N, and S.S. Differential expression was assessed in four pairwise comparisons between the treatments: 1) N.N vs N.S and 2) S.N vs S.S were used to assess the effects of offspring starvation treatment for individuals whose parents experienced the same starvation treatment, whereas 3) N.N vs S.N and 4) N.S vs S.S were used to assess the effects of differing parental starvation treatment on individuals that experienced the same offspring starvation treatment. Significance was based on a Wald Test and shrunken log fold-change values with apeglm (version 1.10.1; Zhu et al. 2019). P-values were adjusted for multiple testing using Benjamini-Hochberg (Benjamini and Hochberg 1995) procedure with a false discovery rate of 0.05. Afterwards, significantly differentially expressed genes (relatively up- or downregulated) were extracted and filtered with a p-adjust of < 0.05. A Venn diagram (Venn.Diagram, version 1.6.20) was used to depict the relationship between the significantly differentially expressed genes for each pairwise comparison. The expression of the significantly differentially expressed genes of each comparison was visualized in a heatmap using normalized counts scaled per gene (scale=“row”) (pheatmap, version 1.0.12).

To examine differential expression of genes with known roles in stress and/or starvation response, we searched specifically for a targeted list of genes within the significantly differentially expressed genes, based on their identity (from the genome annotations). For this, we used the keywords “heat shock protein” (and “hsp”), “cytochrome P450”, “octopamine” and “tyramine” in the putative gene names.

### Pathway-Level Analysis of Differential Expression

We used the KEGG (Kyoto Encyclopedia of Genes and Genomes) database to assign the predicted genes in the *A. rosae* genome to gene pathways using the KEGG Automatic Annotation Server (KAAS; Moriya et al. 2007) (S3). Of the annotated genes for which we had read counts, 65% were assigned to at least one KEGG pathway. A gene set enrichment analysis was then performed on the normalized counts using GAGE (Luo et al. 2009) and pathview (Luo and Brouwer 2013) in R, applying an fasle discovery rate-adjusted *P*-value cut-off of < 0.05 to identify differentially expressed pathways. We derived the non-redundant significant gene set lists, meaning those that did not overlap heavily in their core genes, using esset.grp and a *P*-value cut-off for the overlap between gene sets of 10e^-10^.

Unmapped reads were inspected to check for the differential expression of genes not present in the reference genome. These were extracted from the RSEM-produced BAM file using samtools view (Li et al. 2009) and converted to fastq (using bamtools bamtofastq; Barnett et al. 2011), and *de novo* assembled using Trinity (default settings) (Grabherr et al. 2011). This assembly was done jointly with reads from additional samples (preprint Paul et al. 2021) to optimize coverage. The unmapped reads were mapped back to this reference and expected read counts extracted using eXpress (Roberts et al. 2011). Transcripts with >10,000 mapped reads were identified using BLASTN (default settings, limited to one match per gene and 1e^-1^) against the NCBI nucleotide database. Differential expression of unmapped reads in the larval samples was carried out as for mapped reads (see above).

## Results

### Life-History Traits

Larval starvation in the parental generation had trait- and sex-specific effects on life-history, leading to a prolonged total development time for both sexes and a lower body mass in females (for details see S2). In the offspring generation, the exposure of parents to clerodanoids had no influence on any of the life-history parameters measured, whereas the effect of parental and offspring larval starvation treatment were trait- and sex-specific. Both females (*Chi^2^*_1_ = 11.06, *P* < 0.001) and males (*Chi^2^*_1_ = 29.25, *P* < 0.001) were significantly more likely to have an additional larval instar prior to the eonymph stage if they were starved than if they were not starved (Fig. 2A, B) (analyzed independently of parental starvation treatment). There was an interactive effect of parental and offspring starvation treatment on both female (*Chi^2^*_1_ = 10.37, *P* = 0.001) and male (*Chi^2^*_1_ = 4.89, *P* = 0.027) total development time, such that intra-generational starvation led to a longer development time for both sexes (Fig. 2C, D; S4A, S4B). Larval development of offspring females was only affected by offspring starvation treatment (*Chi^2^*_1_ = 319.40, *P* < 0.001; S4A), while there was an interactive effect of parental and offspring starvation treatment on male larval development time (*Chi^2^*_1_ = 6.60, *P* = 0.010; S4B, S5B). In contrast, for pupal developmental time there was an interactive effect of parental and offspring starvation treatment on females (*Chi^2^*_1_ = 24.57, *P* < 0.001, S4A, S5C), but only a significant effect of offspring starvation treatment on males (*Chi^2^*_1_ = 17.23, *P* < 0.001; S4D). Both parental (*Chi^2^*_1_ = 4.02, *P* = 0.045) and offspring starvation treatment (*Chi^2^*_1_ = 12.95, *P* < 0.001) independently influenced female adult mass, with offspring females having a lower mass if they were starved during development, but those females whose parents were starved had a higher overall mass than when parents had not starved (Fig. 2E). In contrast, male adult mass was significantly affected by the interaction between parental and offspring starvation treatment (*Chi^2^*_1_ = 4.58, *P* = 0.030, Fig. 2F). Male offspring of starved parents that themselves were not starved during development (S.N) had a higher mass than offspring that were starved (S.S) (pairwise comparison: *z* = −3.47, *P* = 0.003, S4B).

### Consumption

Offspring starvation treatment had a significant interactive effect on the relationship between change in mass and initial body mass (relative growth rate) in offspring larvae in both females (*F*_1,60.75_ = 5.96, *P* = 0.017) and males (*F*_1,39.54_ = 7.32, *P* = 0.009; S6a). The change in body mass of starved larvae increased more steeply with an increase in initial larval body mass compared to non-starved larvae for both sexes (Fig. 3A, B), indicating a higher relative growth rate in starved larvae. There was no effect of parental starvation treatment on relative growth rate on adults of both sexes (S6a). The relative consumption rate of females was affected by a significant interaction of initial larval body mass and parental starvation treatment on the area of leaf consumed (*F*_1,62_ = 6.15, *P* = 0.015; S6b), such that the leaf area consumed increased with initial body mass for individuals from non-starved parents but did not change for individuals from starved parents (Fig. 3C). Thus, larger female larvae consumed more than smaller larvae when their parents were not starved but not when their parents were starved. For males, only initial body mass had a significant positive effect on the leaf area consumed (initial body mass: *F*_1,41.45_ = 21.97, *P* < 0.001; S6b) across all treatments (Fig. 3D); i.e. there was no effect of parental or offspring starvation treatment on relative consumption rate. Regarding food conversion efficiency, leaf area consumed and offspring starvation treatment independently influenced change in body mass for both females (leaf area consumed: *F*_1,54.12_ = 7.17, *P* = 0.009; offspring starvation: *F*_1,54.77_ = 11.36, *P* = 0.001) and males (leaf area consumed: *F*_1,44_ = 31.55, *P* < 0.001; offspring starvation: *F*_1,44_ = 6.92, *P* = 0.011; S6c). The food conversion efficiency was higher when larvae were not starved than when they were starved, while under both conditions consuming more leaf material led to a higher body mass increase for larvae (Fig. 3E, F).

**Figure 3:**
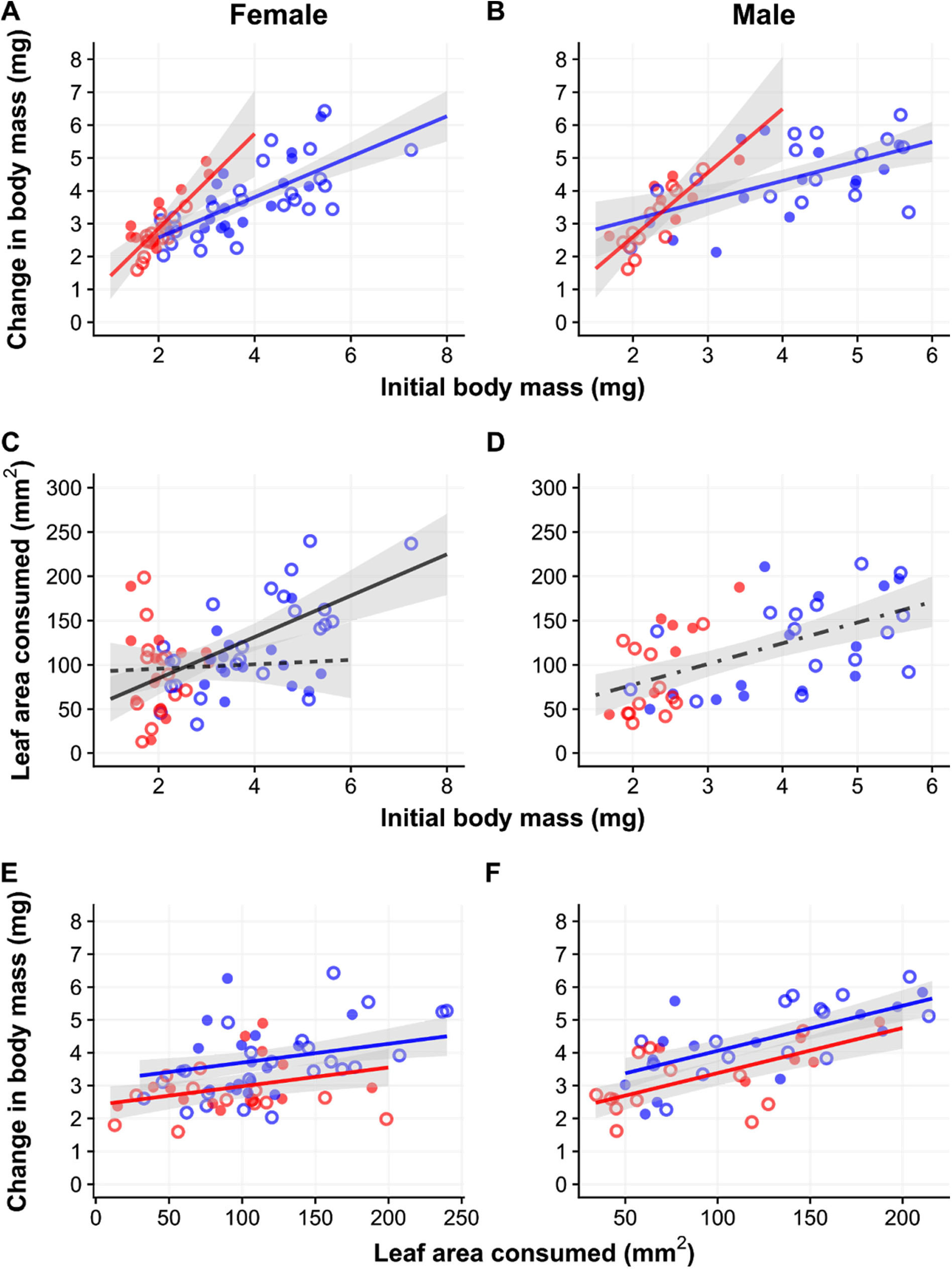
Relationship between initial body mass and change in body mass (relative growth rate: A, B), initial body mass and leaf area consumed (relative consumption rate: C, D), and leaf area consumed and change in body mass (consumption efficiency: E, F) for 3^rd^ instar *Athalia rosae* larvae (offspring generation). Circles represent raw data. Circle color denotes offspring larval starvation treatment (blue=no starvation, red=starvation). Circle type denotes parental larval starvation treatment (open circle=no starvation, filled circle =starvation). Lines depict minimum adequate model predictions with gray shaded regions showing the 95% confidence interval range. Offspring starvation treatment (blue line = no starvation, red line = starvation) had a significant effect on relative growth rate (interactively with initial body mass; A, B) and consumption efficiency (E, F). Parental starvation treatment (solid black line = no starvation, dashed black line = starvation) had a significant effect on relative consumption rate in females (interactively with initial body mass; C).Only initial body mass, but neither parental nor offspring starvation treatment had a significant effect on relative consumption rate in males (dot-dash line in D). Data are plotted separately for females (A, C, E) and males (B, D, F); individuals were sexed on emergence.

### Gene Expression

Sequencing of the 24 individual larvae resulted in a total of 1.4 billion reads with an average of 58 million reads per sample (SE: ±2 million reads) and an average GC content of 43%. Prior to normalization, the average number of mapped reads per sample was 33,675,600, equating to an average of 91% reads per sample aligned to the reference genome (range 85-93%).

Offspring starvation had the strongest effect on gene expression (Figs 4, 5, S7). The two comparisons between starved and non-starved larvae revealed 4727 (N.S vs N.N) and 5089 (S.N vs S.S) significantly differentially expressed genes (log fold-change > 0, adjusted *P* < 0.05) and a large number of these overlapped between the two comparisons (4002). In contrast, there were far fewer significantly differentially expressed genes in the two comparisons in which the larvae had the same starvation treatment, namely 155 (S.N vs N.N) and 75 (N.S vs S.S). In the comparison between starved and non-starved larvae from non-starved parents, there was evidence of regulation in some genes known to be associated with stress response. There was significant downregulation of putative heat shock proteins (eight downregulated and one upregulated) and upregulation of putative cytochrome P450 genes (21 genes upregulated, six downregulated) as well as upregulation of one octopamine and one tyramine. When parental starvation but not offspring starvation treatments differed, there was far less differential expression of such genes with only one or two cytochrome P450 genes being differentially expressed (S8, S9).

**Figure 4:**
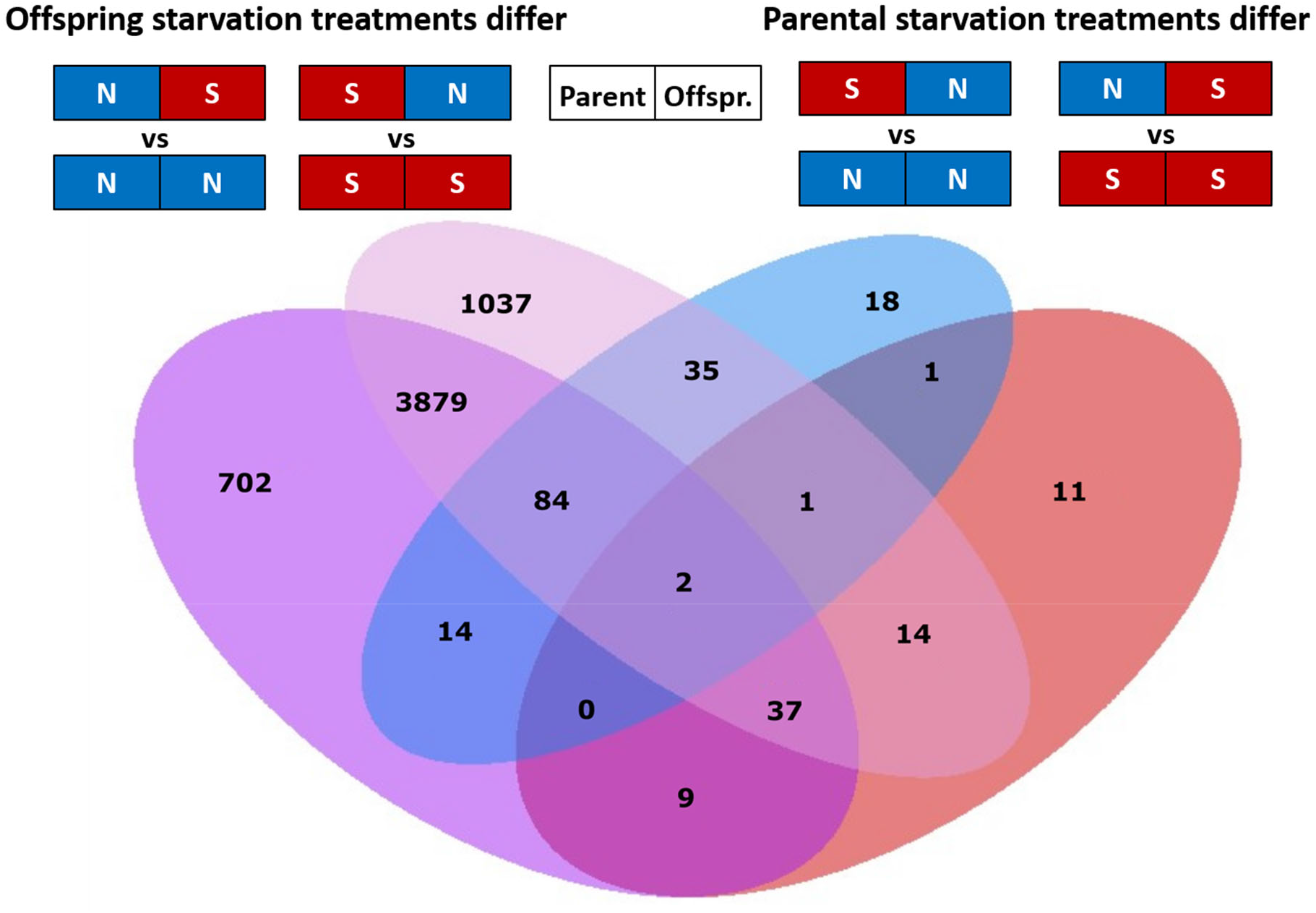
Venn diagram illustrating the number of unique and shared significantly differentially expressed genes (padjust < 0.05) in male offspring larvae of *Athalia rosae* resulting from each of the four pairwise comparisons, (N = no starvation and S = starvation; left box parental, right box offspring treatment).

**Figure 5:**
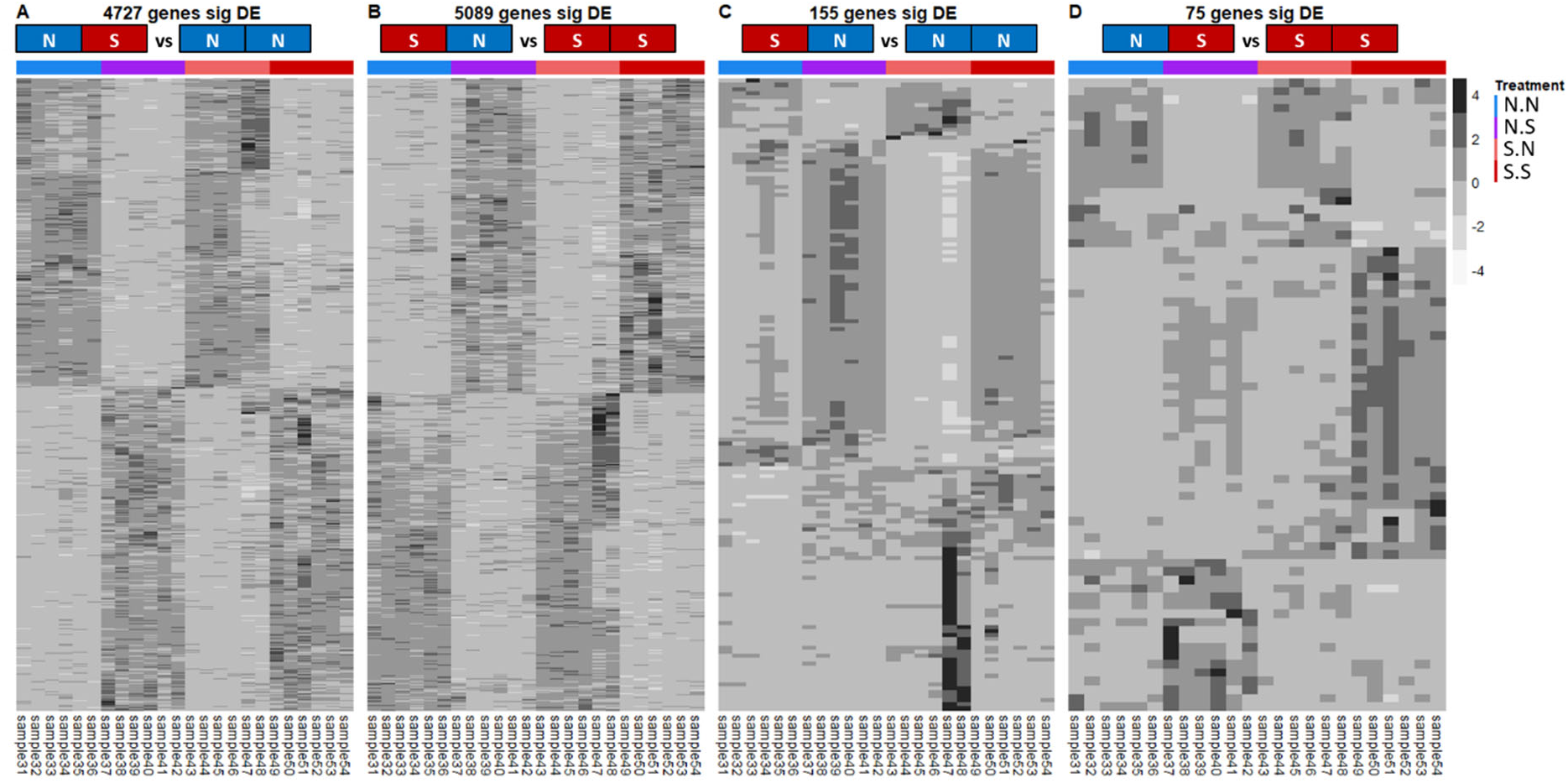
Heatmaps showing the expression (normalized counts) of genes in male offspring larvae of *Athalia rosae* across all samples for those genes that were significantly differentially expressed (padjust < 0.05) when offspring starvation treatment differed (A, B) or parental starvation treatment differed (C, D), (N = no starvation and S = starvation; left box parental, right box offspring treatment). Plotted values are *z*-scores computed from normalized counts post clustering.

While the majority of reads mapped to the reference genome, 247,997,896 reads did not map. We assembled these into 334,717 transcripts; a large proportion of these (267,893 transcripts) were lowly transcribed (<50 reads). Of those genes that we assembled *de novo*, 803 were putatively annotated against the NCBI nt database and of these a large portion were mitochondrial (S10). Differential expression analysis of unmapped reads identified differentially expressed genes between the treatment pairs in a pattern mirroring the genes from the reference [1523 (N.S vs N.N), 1860 (S.N vs S.S), 149 (S.N vs N.N) and 34 (N.S vs S.S)]. Many of these were *A. rosae* genes linked to the mitochondria, particularly in the comparisons between starved and non-starved offspring individuals.

The pathway level analysis with KEGG identified between one and eight pathways that were differentially regulated in our four comparisons (S10). The only pathway that was present in all four comparisons was ko03010, encoding components of the ribosome. Interestingly, expression of ribosomal components was affected not only by differences in offspring starvation treatment but also by differences in parental starvation treatment. The pathway was upregulated in starved larvae when compared to larvae that were not starved (offspring treatment differed) and to larvae that were starved but whose parents were also starved (parental treatment differed), with the reverse trend for non-starved larvae (Table 1; S11). The other significantly differentially expressed KEGG pathways only occurred between individuals that differed in offspring but not parental larval starvation treatments, mirroring the gene expression results. These pathways belong to the four main categories: metabolic breakdown and protein processing (pancreatic secretion, proteasome and protein processing in the endoplasmic reticulum), immune response (phagosome and antigen processing), energy production and central metabolic processes (citrate cycle, glycolysis, thyroid hormone signalling pathway), and ECM-receptor interaction. There was generally a downregulation of these processes in larvae that were starved compared to non-starved individuals.

**Table 1.** Significantly differentially expressed KEGG gene sets/pathways male offspring larvae of *Athalia rosae* that differed in either their own or their parent’s larval starvation regime (N = no starvation and S = two periods of starvation for 24 hours during larval development; left box parental, right box offspring treatment). The combination listed first (on top) is the ‘treatment’ the one listed second (underneath) is the ‘control’ in each comparison, meaning that pathways are up- or downregulated in ‘treatment’ relative to ‘control’. Larvae undergoing starvation (S) were subjected to two periods of starvation for 24 hours during larval development, one in 2^nd^ and one in 4^th^ instar.

| KEGG pathway | Offspring starvation treatment differs |  |  |  | Parental starvation treatment differs |  |  |  |
| --- | --- | --- | --- | --- | --- | --- | --- | --- |
|  | N | S | S | N | S | N | N | S |
|  | vs |  | vs |  | vs |  | vs |  |
|  | N | N | S | S | N | N | S | S |
| Ribosome | ↑ up |  | ↓ down |  | ↓ down |  | ↑ up |  |
| Pancreatic secretion | - |  | ↓ down |  | - |  | - |  |
| Protein processing in endoplasmic reticulum | ↓ down |  | ↑ up |  | - |  | - |  |
| Proteasome | ↓ down |  | ↑ up |  | - |  | - |  |
| Phagosome | - |  | ↑ up |  | - |  | - |  |
| Citrate cycle | ↓ down |  | - |  | - |  | - |  |
| Glycolysis / Gluconeogenesis | ↓ down |  | - |  | - |  | - |  |
| Antigen processing and presentation | ↓ down |  | ↑ up |  | - |  | - |  |
| Protein export | ↓ down |  | - |  | - |  | - |  |
| ECM-receptor interaction | - |  | ↑ up |  | - |  | - |  |
| Thyroid hormone synthesis | - |  | ↑ up |  | - |  | - |  |

## Discussion

We investigated the influence of intra- and intergenerational effects on offspring phenotype in the holometabolous insect *A. rosae*, which realizes distinct niches during its life-cycle. Our results revealed that offspring starvation interacted with parental starvation in a trait-and sex-specific manner, affecting the offspring phenotype. Such trait-specific and sex-specific effects of intergenerational treatments have been shown in previous studies (Zizzari et al. 2016; Le Roy et al. 2017; Wilson et al. 2019; Yin et al. 2019), suggesting that they may be common. Trait- and sex-specificity may result from differential and sex-specific directional selection on traits(Tarka et al. 2018; Yin et al. 2019).

### Effects on Life-History

In our study, larval starvation led to an increased probability of having an additional instar in *A. rosae* larvae of both sexes. Increasing the number of larval instars is a potentially adaptive strategy that allows individuals to recover from starvation via the prolongation of developmental time, e.g. by catch-up growth (Hector and Nakagawa 2012). Such intraspecific variation in the number of larval instars is common in insect species and can occur in response to various biotic and abiotic factors, including food quantity and quality (Esperk et al. 2007). Such variation has also previously been identified in tenthredinid sawfly species (Esperk et al. 2007; Charles and Allan 2000), but has not been described, to our knowledge, in *A. rosae*.

In line with the increased larval instar numbers, offspring of *A. rosae* that were starved had a longer development time in both sexes. There was a trend for faster development times in offspring that experienced a similar environment as their parents, which was at least significant for female pupal development time and may be suggestive of positive effects of matching parental and offspring environmental conditions (Monaghan 2008), i.e. a match-mismatch scenario. Similar positive effects of matching parental and offspring dietary conditions have also been found in a number of species in response to food availability (Hibshman et al. 2016; Raveh et al. 2016). However, other life history data of *A. rosae* did not reveal this match-mismatch pattern, indicating a complex interplay between different intergenerational effects.

Adult mass showed a different pattern than development time, with female offspring having a higher adult mass when their parents were starved as larvae irrespective of their own starvation conditions, indicative of enhanced parental provisioning to offspring of starved parents, i.e. parental buffering. Similarly, females of the vinegar fly, *Drosophila melanogaster*, reared on poor food had larger offspring than females reared on standard food (Valtonen et al. 2012). Life-history theory predicts that under hostile conditions there is a shift towards fewer but better provisioned offspring (Roff 1992), although we did not examine the number of offspring produced here. As neither larval developmental time for female *A. rosae* nor their consumption were affected by the parental starvation treatment, other physiological effects (discussed below for gene expression) may have mediated this kind of buffering. In contrast to females, only male offspring of starved parents differed significantly in adult mass, with starved sons weighing less. Thus, investment in body mass may be particularly low in males under repeatedly poor environmental conditions across generations, indicating “parental suffering” rather than buffering (i.e. negative parental effects). The differential investment in sexes in dependence of the parental vs. offspring starvation may be related to the haplodiploidy of this sawfly species. For example, larger eggs are usually fertilized to become females while smaller eggs remain unfertilized and become males in the haplodiploid thrip, *Pezothrips kellyanus*, and such differences could increase under parental starvation conditions (Katlav et al., 2021). Overall, there was no clear indication of silver spoon effects for any of the measured life-history traits, but in match-mismatch designs such effects cannot be ruled out (Engqvist and Reinhold 2016).

### Effects on Consumption

Consumption assays revealed a different pattern than expected from the body mass patterns. Considering the larger body mass of female offspring from starved parents, we would have expected such individuals to have a higher growth rate, consume more leaf material, and/or have a higher consumption efficiency. Our results showed that parental starvation had no effect on any consumption trait except on relative consumption rate in females, where offspring of non-starved parents consumed more leaf material than that of starved parents, contrary to our expectation. Additionally, offspring that experienced starvation exhibited a steeper relative growth rate than non-starved larvae, suggesting compensatory growth (Hector and Nakagawa 2012). Combining this compensatory growth with a longer development period potentially allows starved individuals to become larger and to overcome size-based selection, which can be an important determinant of individual fitness for many insect species (Beukeboom 2018). While such compensatory growth has been seen in many species, it can also pose costs to individuals (Arendt et al. 2001; Dmitriew and Rowe 2007; Auer et al. 2010). We found that individuals that were starved had a lower food conversion efficiency than non-starved individuals, which may suggest a physiological cost of compensatory growth. Adjustments in consumption patterns to different starvation regimes have been revealed to be expressed in the next generation (McCue et al. 2017). However, patterns may differ depending on the developmental stage in which an individual is facing starvation, which needs to be further explored in *A. rosae*. Since we conducted the consumption assays for all larvae at the same developmental stage, we were able to disentangle the effect of starvation from effects due to differences in ontogeny (Nicieza and Álvarez 2009), in contrast to earlier studies on this sawfly (Paul et al. 2019).

### Effects of Parental Reproductive Environment

Unlike starvation and in contrast to our expectation, clerodanoid uptake, which is known to enhance mate attractiveness in *A. rosae* (Amano et al. 1999; preprint Paul and Müller 2021), had no effect on the measured traits in offspring of both sexes. This is surprising, because partner attractiveness can affect investment in offspring or/and offspring traits (Robart and Sinervo 2019), sometimes having multigenerational consequences (Gilbert et al. 2012). In other species, mating with more attractive partners can lead to direct effects for the partner, but not necessarily for the offspring. For example, in the field cricket, *Gryllus firmus*, mating with more attractive males led to a higher number of eggs laid by females but offspring did not show any fitness benefits (Kelly and Adam-Granger 2020). In *A. rosae*, parental clerodanoid exposure may affect other offspring traits, such as immunity (Bozov et al. 2015), lifespan (Zanchi et al. 2021), or traits exhibited in adulthood, e.g. mating success (Amano et al. 1999; preprint Paul and Müller 2021), that were not measured in our study.

### Effects on Gene Expression

Similar to the findings for life-history and consumption, transcriptome analysis of male larvae of the offspring generation also revealed stronger intra-than intergenerational effects of larval starvation on gene expression, in line with our hypothesis. When offspring larvae were starved, about half of the genes were differentially expressed compared to non-starved larvae. As typical stress response indicators, genes putatively encoding heat shock proteins and cytochrome P450s were differentially expressed, being both up- and down-regulated in *A. rosae* in response to starvation. Regulation of heat shock proteins in response to starvation has been reported elsewhere, including in larvae of Lepidoptera and Hymenoptera (Farahani et al. 2020; Wang et al. 2012). These proteins act to stabilize and protect other proteins in the face of both abiotic and biotic stresses (Sørensen et al. 2003; Farahani et al. 2020). Cytochrome P450s are best known for their role in metabolism of xenobiotics by insects (Feyereisen 2012). However, they were also found to be more highly expressed in *D. melanogaster* flies selected for starvation resistance compared to controls (Doroszuk et al. 2012). In addition, genes putatively encoding the monoamines octopamine and tyramine were upregulated in starved larvae of *A. rosae*, both of which play a key role in modulating behavioral and physiological processes in invertebrates and can be involved in starvation resistance (Li et al. 2016).

Furthermore, starvation of the offspring led to differential gene expression of pathways involved in metabolism, including pancreatic secretion, which was also found in starved larvae of the cotton bollworm, *Helicoverpa armigera* (Lepidoptera: Noctuidae) (Jiang et al. 2019). In starved larvae of *A. rosae*, differential regulation of metabolic pathways may explain why these larvae showed an overall lower food conversion efficiency compared to non-starved larvae. Moreover, expression of metabolic genes likely changes with the duration of starvation. For example, in *D. melanogaster* a 16 h starvation period led to downregulation particularly of the endopeptidase gene family (Moskalev et al. 2015), but extended periods of starvation may induce an activation of proteolysis genes. However, little is known about long-term effects across generations, on gene expression patterns in response to starvation. Interestingly, many of the genes differentially expressed in starved vs non-starved individuals of *A. rosae* linked to the mitochondria, reflecting wholescale disruption of homeostasis caused by starvation (Gilbert 2012).

Parental starvation also caused differential expression in about 1 % of genes in *A. rosae* offspring, indicating that subtle intergenerational imprints on gene expression can occur. When only parental, but not offspring, treatment differed, offspring from starved parents displayed a downregulation of the ribosomal pathway compared to offspring of non-starved parents. In contrast, when only offspring treatment differed, larval starvation caused a comparative enrichment of genes in this pathway. These results indicate that larval starvation can have significant direct effects on the regulation of ribosomal proteins, but that these effects may be buffered to some degree via parental starvation. Starved larvae from non-starved parents may have been more physiologically stressed than starved larvae from starved parents, as indicated by the enrichment of putative ribosomal pathway genes. Differential regulation of genes in this pathway has been observed in a wide variety of taxa in response to thermal stress (Paraskevopoulou et al. 2020; Schwanz et al. 2020; Srikanth et al. 2020), demonstrating the broad role of this pathway in stress response. Importantly, the differential expression of genes involved in stress and other physiological and metabolic responses may have energetic costs, which may contribute to a prolonged development time of starved larvae in *A. rosae*.

Alteration in genes encoding ribosomal proteins may also be an important mechanism mediating intergenerational effects of starvation (Aldrich and Maggert 2015; Bughio and Maggert 2019). In *Caenorhabditis elegans* starvation led to the generation of small RNAs, and small RNA-induced gene silencing persisted over up to three generations (Rechavi et al. 2014). Such mechanisms may also explain the intergenerational changes in gene expression observed in *A. rosae* in the present study; future work comparing small RNA expression in relation to starvation could address this.

## Conclusions

In summary, our findings highlight that intragenerational starvation effects were somewhat stronger than the intergenerational effects of starvation across life-history, consumption, and gene expression patterns. We found some evidence for parental effects including match-mismatch and parental buffering or parental suffering, while there was no clear evidence for silver spoon effects. The parental reproductive environment left no signature on the measured offspring traits. Due to different trajectories and environments experienced during different stages, distinct niches are realized, which are expressed in diverse phenotypes. Our results suggest that periods of starvation may make individuals more robust when facing another food shortage in the next generation and may, in the case of herbivorous pest insects, lead to enhanced damage of crops.

## Acknowledgments

This study was funded by the German Research Foundation (DFG) as part of the SFB TRR 212 (NC^3^), project number 396777467 (granted to CM). We also thank University of Potsdam AG Genetics and AG Evolutionary Adaptive Genomics for use of computational resources and to Dr Tobias Busche and Katrin Lehmann for assistance with RNA extraction.

## Data availability

All data and code of this manuscript will be deposited online on Dryad (DOI https://doi.org/10.5061/dryad.73n5tb2x0).

Raw Reads: SRA ID = PRJNA716060. BioSample accession numbers = SAMN18393262-SAMN18393285

## Author contributions

Conceptualization and funding acquisition: CM; methods development/experimental design: SCP, CM; data collection: SCP, data validation and analysis: life history data: SCP, consumption assay data: SCP, PS, gene expression data: SCP, ABD; data visualization: SCP, PS; writing original draft: SCP, CM, PS; reviewing and editing: SCP, PS, ABD, CM.

## Supplemental Material

S1. Full Breeding Design of Experimental Animals

S2. Effects of Starvation on Life-History Traits of Parental Generation

S3. KEGG Term Analysis

S4. Results of Posthoc Analyses for Offspring Generation

S5: Influence of Parental and Offspring Larval Starvation Treatments on Developmental Times of *Athalia rosae* in Offspring Generation

S6. Effects of Predictor Variables and Their Interactions on Consumption Traits.

S7. Percentage of Differentially Expressed Genes

S8. Numbers of Significantly Up- or Downregulated Genes

S9. Mapped reads, DE output.

S10. Unmapped reads, DE output.

S11. KEGG pathway results.

## Notes

### Competing Interest Statement

The authors have declared no competing interest.

### Summary of Updates

Methods and results section updated for further clarification and entire text revised. Figures 1-3 revised. Supplement files updated.

