## Supplement Material for "Intergenerational Effects of Early Life Starvation on Life-History, Consumption, and Transcriptome of a Holometabolous Insect"

### S1. Full Breeding Design of Experimental Animals

Adults of *A. rosae* (F0) were collected in May 2019 from meadows adjacent to farmland in Germany at two locations (population A: 52°02'48.0"N 8°29'17.7"E, population B: 52°03'54.9"N 8°32'22.2"E). Individuals of each population were split between two cages (total of 4 cages) and maintained for one generation at room temperature and 16 h: 8 h light:dark on potted plants of white mustard (*Sinapis alba*) and Chinese cabbage (*Brassica rapa* var. *pekinensis*). F1 adults were used to set up a total of six cages with males and females from different populations in four cages and two cages with virgin females only (Fig. S1). Larvae of the final instar, the eonymphs, were placed into individual cups containing soil for pupation. Emerged adults were kept individually in Petri dishes and provided with a honey:water mixture (1:50). For mating, pairs of non-sib F2 females and males (N = 20 per treatment level) were placed together and allowed to mate at least once. Afterwards, mated as well as virgin females were placed individually into boxes (25 x 15 x 10 cm) with a middle-aged leaf of 6-8 week old cabbage plants supplied with water for oviposition and a honey:water mixture in a climate chamber (20 °C:16 °C, 16 h: 8 h light:dark, 70% r.h.). The boxes were checked and topped up with honey water daily, and an additional leaf was added if the first leaf showed signs of wilting. Females were removed from the boxes after one week and their offspring used to set up the experimental generations (Fig. S1).

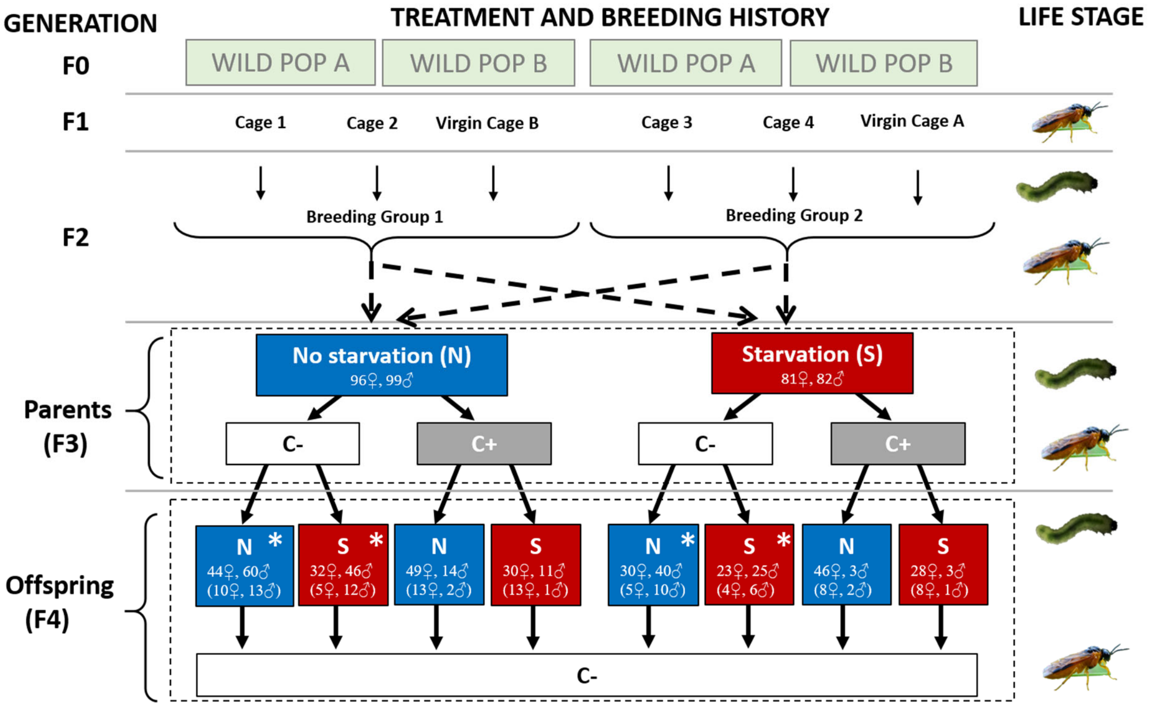


**Figure S1:** Full experimental design showing outline of breeding structure and experimental treatments from F0 to F4 generation in *Athalia rosae*; * indicates individuals taken for RNASeq analysis. Sample sizes split by sex are given in boxes for F3 and F4 generation. Numbers in brackets refer to sample sizes for consumption assay (please note that individuals were pooled across C+ and C- treatments).

### S2. Effects of Starvation on Life-History Traits of Parental Generation

#### *Methods*

Variation in adult mass, larval, pupal, and total developmental time of individuals of the parental generation were assessed using a linear mixed model (LMM, package: lme4; ; version 1.1-23), in which starvation treatment (N, S), sex, and their interaction were the fixed effects and parental pair was included as a random effect to control for possible parental effects (non-independence of sibling larvae taken from the same breeding box).

#### *Results*

Starvation during development had sex-specific effects on life-history traits in the parental generation. When larvae were starved during development, their developmental time was prolonged compared to non-starved individuals and this effect was larger in males than females (starvation * sex: $Chi^2_1 = 7.75$, $P = 0.005$; pairwise comparisons males: Estimate ± SE= 3.92 ± 0.26, $z = 15.35$, $P < 0.001$; females: 2.91 + 0.26, $z = 11.32$, $P < 0.001$; S1 Table 1A, S1 Fig. 1A, B). When broken down into larval and pupal development times, starvation prolonged larval development time equally for both males and females (starvation * sex: n.s.; starvation: $Chi^2_1 = 325.54$, $P < 0.001$, S1 Fig. 1C, D), but only led to a prolonged pupal development time in males (starvation * sex: $Chi^2_1 = 7.05$, $P = 0.007$; pairwise comparisons: males $z = 5.57$, $P < 0.001$; females $z = 1.78$, $P = 0.280$; S1 Table 1B; S1 Fig. 1E, F). In contrast, only female but not male adult body mass was reduced due to larval starvation (starvation * sex: $Chi^2_1 = 10.09$, $P = 0.001$; pairwise comparisons: males $z = 1.22$, $P = 0.610$; females $z = -3.28$, $P = 0.005$; S1 Table 1C; S1 Fig. 1G, H).

### ***Discussion***

Within the parental generation, females had a lower body mass when larvae had been exposed to starvation, while males were able to achieve a similar adult mass despite early life starvation. The result for males may partly be explained by the fact that their development was comparatively more prolonged by larval starvation than that of females. With a particularly long pupal developmental time they could then catch-up, reaching a similar adult body mass as non-starved males. Such catch-up growth occurs in a wide range of taxa (Hector and Nakagawa 2012). In contrast, in an earlier study with *A. rosae*, no catch-up growth was found in individuals when larvae were exposed to slightly different starvation regimes, i.e., starvation bouts of 24 h either every 3 or 4 days during the larval development; both females and males showed prolonged development times and reached a lower adult body mass than well-fed individuals under these conditions (Paul et al. 2019). Thus, the ability to catch-up may highly depend on the strength of the stress applied.

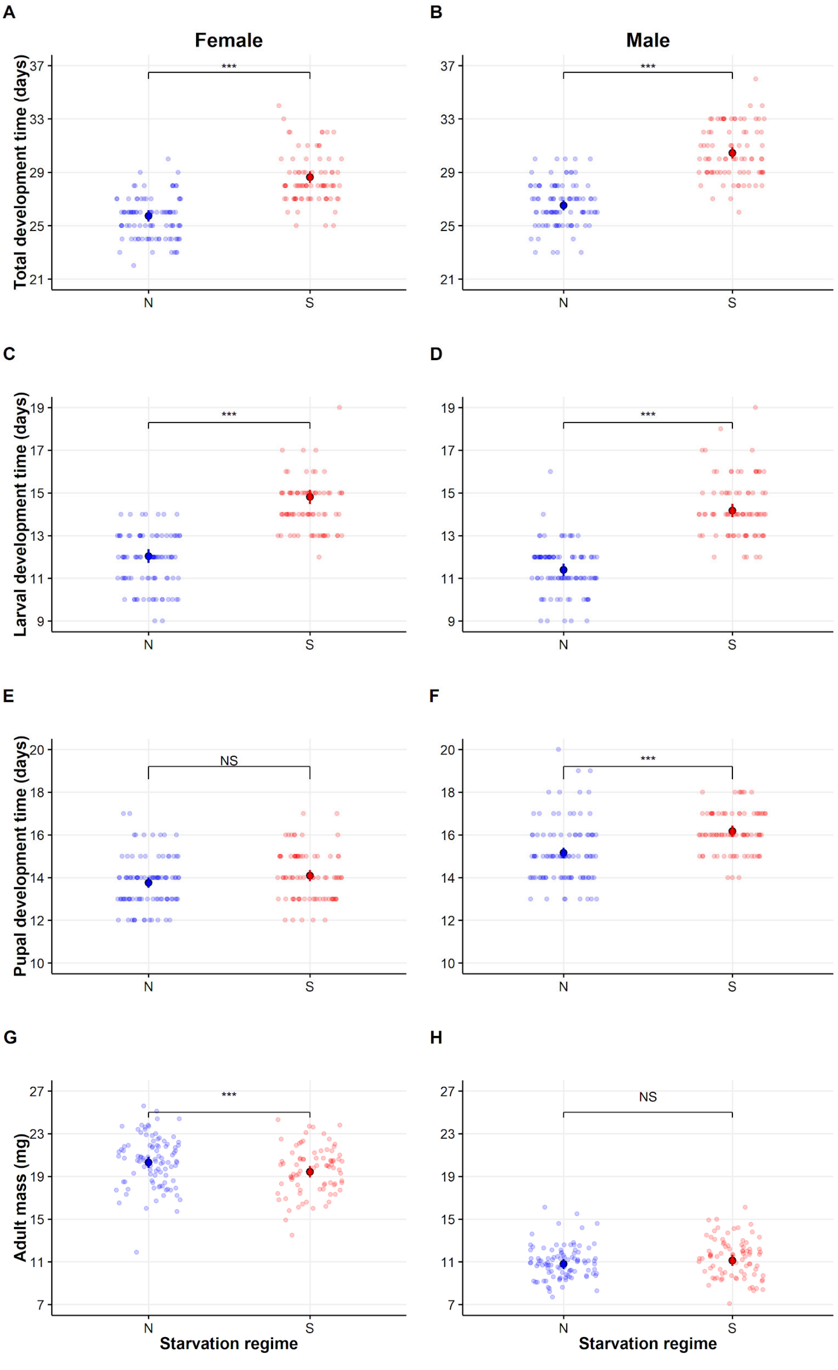


**S2 Figure 1**. Total, larval, and pupal development time as well as adult mass of non-starved (N) and starved (S) *Athalia rosae* (parental generation). Points are model predictions with associated confidence intervals and colors of points (N-blue, S-red) correspond to starvation treatment. Significant differences between N and S treatment are indicated by *** (Tukey's HSD, $\alpha = 0.05$). Raw data is plotted in transparent colours in the background.

**S2 Table 1**. Result of posthoc analyses (Tukey's HSD, $\alpha = 0.05$), obtained using the package ‘multcomp’, for (a) total development time, (b) pupal development time, and (c) adult mass of the parental generation of *Athalia rosae*. S and N indicate starvation and non-starvation treatment, respectively. Significant differences are highlighted in bold.

| Trait | Pairwise comparison | Estimate | SE | *z* value | *Pr*(>/z/) |
|---|---|---|---|---|---|
| A) Total development time | S♀ - N♀ | 2.909 | 0.257 | 11.32 | **<0.001** |
| | N♂ - N♀ | 0.799 | 0.270 | 2.96 | **0.016** |
| | S♂ - N♀ | 4.723 | 0.282 | 16.72 | **<0.001** |
| | N♂ - S♀ | -2.111 | 0.279 | -7.57 | **<0.001** |
| | S♂ - S♀ | 1.814 | 0.292 | 6.21 | **<0.001** |
| | S♂ - N♂ | 3.924 | 0.256 | 15.35 | **<0.001** |
| B) Pupal development time | S♀ - N♀ | 0.326 | 0.183 | 1.78 | 0.28 |
| | N♂ - N♀ | 1.401 | 0.174 | 8.06 | **< 0.001** |
| | S♂ - N♀ | 2.410 | 0.182 | 13.21 | **<0.001** |
| | N♂ - S♀ | 1.075 | 0.182 | 5.92 | **<0.001** |
| | S♂ - S♀ | 2.084 | 0.190 | 10.97 | **<0.001** |
| | S♂ - N♂ | 1.009 | 0.181 | 5.57 | **<0.001** |
| C) Adult mass | S♀ - N♀ | -0.0009 | 0.0003 | -3.28 | **0.006** |
| | N♂ - N♀ | -0.010 | 0.0003 | -31.33 | **< 0.001** |
| | S♂ - N♀ | -0.009 | 0.0003 | -28.97 | **< 0.001** |
| | N♂ - S♀ | -0.009 | 0.0003 | -27.66 | **< 0.001** |
| | S♂ - S♀ | -0.008 | 0.0003 | -25.46 | **< 0.001** |
| | S♂ - N♂ | 0.001 | 0.0003 | 1.22 | 0.61 |

***References***


Hector, K. L., and S. Nakagawa. 2012. Quantitative analysis of compensatory and catch-up growth in diverse taxa. Journal of Animal Ecology 81:583-593.

Paul, S. C., R. Putra, and C. Müller. 2019. Early life starvation has stronger intra-generational than transgenerational effects on key life-history traits and consumption measures in a sawfly. Plos One 14:e0226519.

### S3. KEGG Term Analysis

For characterization of the differential expression of pathways in the different treatments, KEGG (Kyoto Encyclopedia of Genes and Genomes) terms were allocated to the gene expression data using the KEGG Automatic Annotation Server (KAAS; Moriya et al. 2007). A bi-directional best hit KAAS was run using the predicted gene sequences from the annotated *A. rosae* genome, with 40 different insect species selected for reference (dme, mde, lcq, aga, aag, aalb, cqu, ame, bim, bter, ccal, obb, soc, mpha, aec, acep, pbar, vem, hst, dqu, cfo, lhu, pgc, obo, pcf, nvi, mdl, tca, dpa, atd, nvl, bmor, bman, dpl, pmac, prap, haw, tnl, pxy, api). In several cases (< 30), in which multiple KEGG terms were assigned to the same gene, read counts were duplicated for each individual KEGG term, and when multiple genes were given the same KEGG term, read counts were summed across all genes that matched to each term.

#### *Reference*


Moriya, Y., M. Itoh, S. Okuda, A. C. Yoshizawa, and M. Kanehisa. 2007. KAAS: an automatic genome annotation and pathway reconstruction server. Nucleic Acids Research 35:W182-W185.

**S4. Results of Posthoc Analyses for Offspring Generation**

**A**. Result of posthoc analyses (Tukey's HSD, $\alpha = 0.05$), obtained using the package 'multcomp', for total development time, and pupal development time of females of *Athalia rosae* in the offspring generation. Significant differences are highlighted in bold. For the treatment notation, the first letter refers to the parental starvation treatment while the second letter (separated by ".") refers to the offspring starvation treatment, with S indicating starvation and N indicating non-starvation treatment.

| Trait | Pairwise comparison | Estimate | SE | *z* value | Pr(>/z/) |
|---|---|---|---|---|---|
| Total development time | S.N - N.N | 0.17 | 0.21 | 0.81 | 0.84 |
| | N.S - N.N | 3.05 | 0.14 | 21.37 | **<0.001** |
| | S.S - N.N | 2.52 | 0.22 | 11.52 | **<0.001** |
| | N.S - S.N | 2.88 | 0.22 | 13.22 | **<0.001** |
| | S.S - S.N | 2.36 | 0.16 | 14.93 | **<0.001** |
| | S.S - N.S | -0.52 | 0.23 | -2.28 | 0.1 |
| Pupal development time | S.N - N.N | 0.42 | 0.13 | 3.16 | **0.008** |
| | N.S - N.N | 0.74 | 0.11 | 6.71 | **< 0.001** |
| | S.S - N.N | 0.33 | 0.14 | 2.26 | 0.11 |
| | N.S - S.N | 0.32 | 0.14 | 2.26 | 0.11 |
| | S.S - S.N | -0.09 | 0.12 | -0.77 | 0.86 |
| | S.S - N.S | -0.42 | 0.15 | -2.72 | **0.03** |

**B**. Result of posthoc analyses for total development time, larval development time, and adult mass of males of *Athalia rosae* in the offspring generation. Significant differences are highlighted in bold. For the treatment notation, see legend of Table A.

| Trait | Pairwise comparison | Estimate | SE | *z* value | *Pr*(>/z/) |
|---|---|---|---|---|---|
| Total development time | S.N - N.N | 0.44 | 0.31 | 1.42 | 0.48 |
| | N.S - N.N | 3.39 | 0.23 | 14.69 | **<0.001** |
| | S.S - N.N | 2.96 | 0.33 | 8.86 | **<0.001** |
| | N.S - S.N | 2.96 | 0.32 | 9.36 | **<0.001** |
| | S.S - S.N | 2.53 | 0.32 | 7.94 | **<0.001** |
| | S.S - N.S | -0.43 | 0.34 | -1.26 | 0.59 |
| Larval development time | S.N - N.N | 0.24 | 0.22 | 1.12 | 0.67 |
| | N.S - N.N | 2.88 | 0.17 | 16.53 | **<0.001** |
| | S.S - N.N | 2.36 | 0.24 | 9.77 | **<0.001** |
| | N.S - S.N | 2.63 | 0.23 | 11.64 | **<0.001** |

| | | | | | |
|---|---|---|---|---|---|
| | S.S - S.N | 2.12 | 0.24 | 8.82 | **<0.001** |
| | S.S - N.S | -0.52 | 0.25 | -2.08 | 0.16 |
| Adult mass | S.N - N.N | 0.0007 | 0.0004 | 1.77 | 0.28 |
| | N.S - N.N | -0.0002 | 0.0002 | -1.01 | 0.73 |
| | S.S - N.N | -0.0005 | 0.0004 | -1.24 | 0.59 |
| | N.S - S.N | -0.0009 | 0.0004 | -2.38 | 0.08 |
| | S.S - S.N | -0.0011 | 0.0003 | -3.47 | **0.002** |
| | S.S - N.S | -0.0002 | 0.0004 | -0.61 | 0.93 |

**S5: Influence of Offspring and Parental Larval Starvation Treatments on Developmental Times of *Athalia rosae* in Offspring Generation**

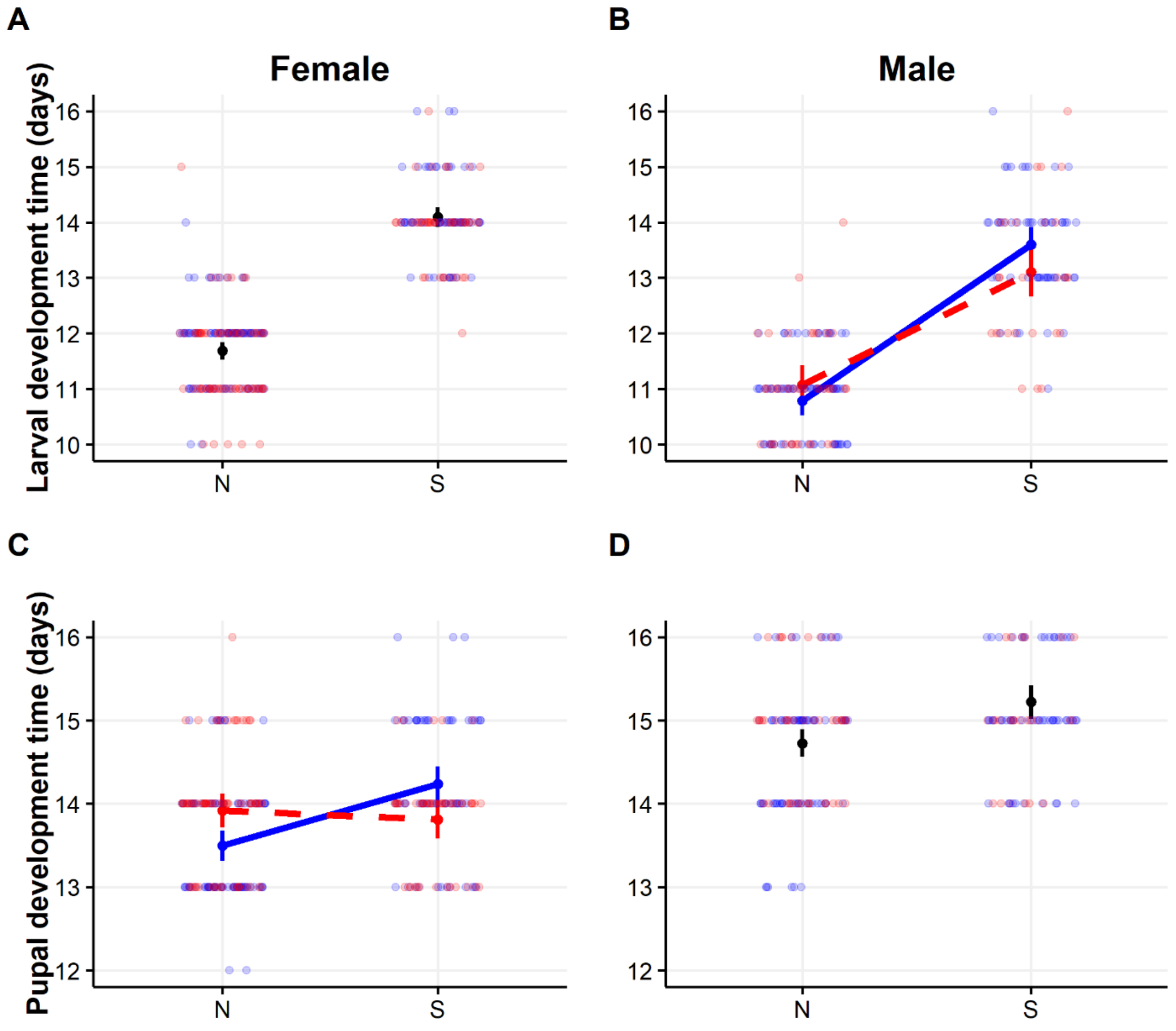


**S5**. Influence of offspring larval starvation treatments (N = no starvation, S = starvation) and parental larval starvation treatments (blue solid line = no parental starvation, red dashed line = parental starvation) on offspring larval development time (A, B) and pupal development time (C, D) of *Athalia rosae* in F4 generation. Data are plotted separately for females (A, C) and males (B, D); individuals were sexed on emergence. Points are model predictions with associated confidence intervals and colors of points and lines correspond to parental starvation treatment. Raw data is plotted in the background in transparent colors (blue circles = no parental starvation, red circles = parental starvation). Note that there was no significant effect of parental starvation treatment for female larval development time and male pupal development time.

**S6. Effects of Predictor Variables and Their Interactions on Consumption Traits**

Values from lmm (using conditional *F*-test with Satterthwaite approximation). Significant effects (P< 0.05) are highlighted in bold.

| Trait | Predictor variables | Female | | | Male | | |
|---|---|---|---|---|---|---|---|
| | | *F* | *df* | *P* | *F* | *df* | *P* |
| a) Relative growth rate | Initial_Body_mass*Parental_Starvation_Treatment*Offspring_Starvation_Treatment | 0.21 | 1, 57.9 | 0.648 | 1.84 | 1, 33.4 | 0.184 |
| | Parental_Starvation_Treatment*Offspring_Starvation_Treatment | 0.26 | 1, 60.0 | 0.613 | 1.34 | 1, 39.8 | 0.254 |
| | Initial_Body_mass*Parental_Starvation_Treatment | 1.49 | 1, 43.2 | 0.229 | 0.003 | 1, 35.6 | 0.952 |
| | Initial_Body_mass*Offspring_Starvation_Treatment | **5.96** | **1, 60.8** | **0.017** | **7.32** | **1, 39.5** | **0.009** |
| | Initial_Body_mass | - | - | - | - | - | - |
| | Parental_Starvation_Treatment | 3.59 | 1, 61.0 | 0.063 | 0.01 | 1, 17.4 | 0.911 |
| | Offspring_Starvation_Treatment | - | - | - | - | - | - |
| b) Relative consumption rate | Initial_Body_mass*Parental_Starvation_Treatment*Offspring_Starvation_Treatment | 0.35 | 1, 58.0 | 0.886 | 0.32 | 1, 34.0 | 0.574 |
| | Parental_Starvation_Treatment*Offspring_Starvation_Treatment | 0.40 | 1, 59.0 | 0.528 | 2.63 | 1, 40.8 | 0.112 |
| | Initial_Body_mass*Parental_Starvation_Treatment | **6.15** | **1, 62.0** | **0.015** | 2.69 | 1, 34.0 | 0.110 |
| | Initial_Body_mass*Offspring_Starvation_Treatment | 1.07 | 1, 60.0 | 0.305 | 2.03 | 1, 38.4 | 0.162 |
| | Initial_Body_mass | - | - | - | **21.97** | **1, 41.4** | **< 0.001** |
| | Parental_Starvation_Treatment | - | - | - | 0.48 | 1, 14.2 | 0.496 |
| | Offspring_Starvation_Treatment | 0.13 | 1, 61.0 | 0.713 | 1.57 | 1, 39.2 | 0.217 |
| c) Food conversion efficiency | Leaf_area_consumed*Parental_Starvation_Treatment*Offspring_Starvation_Treatment | 0.88 | 1, 58.0 | 0.351 | 0.38 | 1, 38.1 | 0.537 |
| | Parental_Starvation_Treatment*Offspring_Starvation_Treatment | 0.17 | 1, 52.8 | 0.676 | 0.84 | 1, 39.5 | 0.364 |
| | Leaf_area_consumed*Parental_Starvation_Treatment | 0.004 | 1, 45.6 | 0.947 | 0.04 | 1, 39.9 | 0.829 |
| | Leaf_area_consumed*Offspring_Starvation_Treatment | 2.72 | 1, 59.9 | 0.104 | 1.07 | 1, 40.6 | 0.306 |

| | | | | | | |
|---|---|---|---|---|---|---|
| Leaf_area_consumed | **7.17** | **1, 54.1** | **0.009** | **31.55** | **1, 44.0** | **< 0.001** |
| Parental_Starvation_Treatment | 1.66 | 1, 25.0 | 0.209 | 0.02 | 1, 42.9 | 0.875 |
| Offspring_Starvation_Treatment | **11.36** | **1, 54.7** | **0.001** | **6.92** | **1, 44.0** | **0.011** |

**S7. Percentage of Differentially Expressed Genes**

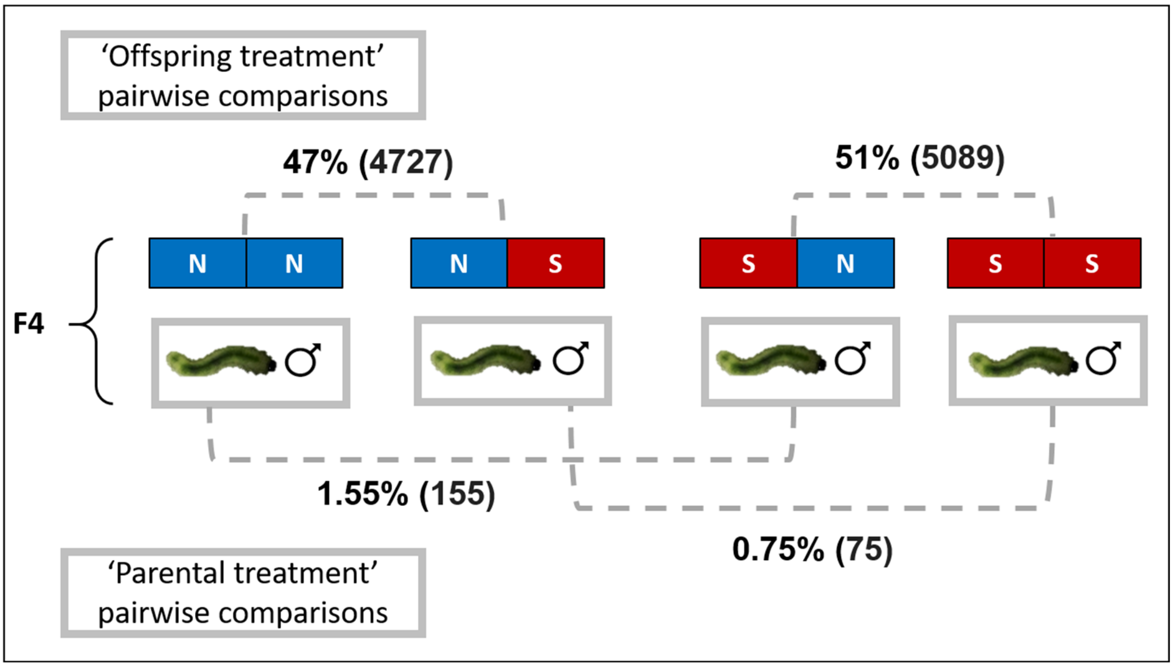


**S7.** Percentage (and number) of genes that were differentially expressed in four pairwise comparisons between male offspring larvae of *Athalia rosae* that differed in either their own or their parent's larval starvation regime (N = no starvation and S = starvation, left box parental, right box offspring treatment), each out of a total of 10,024 genes with non-zero total read count.

**S8. Numbers of Significantly Differentially Expressed Genes.**

| Putative gene ID | Offspring starvation differs | | | Parental starvation differs | |
|---|---|---|---|---|---|
| | N S | S N | | S N | N S |
| | vs | vs | | vs | vs |
| | N N | S S | | N N | S S |
| **heatshock proteins (hsp)** | **1 ↑ - 8 ↓** | **0 ↑ - 0 ↓** | | **0 ↑ - 0 ↓** | **0 ↑ - 0 ↓** |
| **cytochrome P450** | **21 ↑ - 6 ↓** | **2 ↑ - 0 ↓** | | **2 ↑ - 0 ↓** | **0 ↑ - 1 ↓** |
| **octopamine** | **2 ↑ - 0 ↓** | **0 ↑ - 0 ↓** | | **0 ↑ - 0 ↓** | **0 ↑ - 0 ↓** |
| **tyramine** | **1 ↑ - 0 ↓** | **0 ↑ - 0 ↓** | | **0 ↑ - 0 ↓** | **0 ↑ - 0 ↓** |
